# Assessing metadata and curation quality: a case study from the development of a third-party curation service at Springer Nature

**DOI:** 10.1101/530691

**Authors:** Rebecca Grant, Graham Smith, Iain Hrynaszkiewicz

## Abstract

Since 2017, the publisher Springer Nature has provided an optional Research Data Support service to help researchers deposit and curate data that support their peer-reviewed publications. This service builds on a Research Data Helpdesk, which since 2016 has provided support to authors and editors who need advice on the options available for sharing their research data. In this paper we describe a short project which aimed to facilitate an objective assessment of metadata quality, undertaken during the development of a third-party curation service for researchers (Research Data Support). We provide details on the single-blind user-testing which was undertaken, and the results gathered during this experiment. We also briefly describe the curation services which have been developed and introduced following an initial period of testing and piloting. This paper will be presented at the International Digital Curation Conference 2019, and has been submitted to the International Journal of Digital curation.

## Introduction

In 2016 the publisher Springer Nature introduced standard research data policies for its journals (Hrynaszkiewicz et al, 2017) with the aim of encouraging each journal to select the policy which is most appropriate for its discipline and its community. To date, more than 1500 Springer Nature journals have selected a standard policy. Four policies are offered, which include consistent features such as data citation, data availability statements, data deposition in repositories and data peer review.

The introduction of standard data policies by publishers is in response to growing demand from research funding agencies for data sharing. There is also evidence that the research data policies of journals and publishers, historically, have lacked standards and were in need of harmonisation (Naughton & Kernohan, 2016).

Since the introduction of standard data policies by Springer Nature, other academic publishers have begun to introduce similar policies, including Taylor & Francis, Elsevier, Wiley and BMJ. A global Research Data Alliance Interest Group^1^ has also developed a master policy framework with the aim of harmonising data policies across all publishers, and this was published in draft format in 2018. With the on-going introduction of data policies by publishers, as well as funding agencies and institutions, researchers are increasingly compelled to share the data which underpins their published articles.

In 2016, as the standard data policies began rolling out across journals, Springer Nature introduced a Research Data Helpdesk to provide support to researchers, authors and academic editors who need to comply with or implement journal data policies. The Helpdesk also provides email-based support to researchers and editors regarding many aspects of data sharing and publication. Although all aspects of research data are covered by the Helpdesk including data preservation, licensing and publishing, an analysis of the Helpdesk queries in 2017 found that the majority related to data policies and policy compliance. Researchers also requested information regarding the deposit of data in repositories, and how data availability statements should be drafted, which also reflect the requirements of their journal’s data policies (Astell, Hrynaszkiewicz, Grant, Smith & Salter, 2017).

In 2017 Springer Nature undertook a large-scale survey of researchers that received nearly 8000 responses and aimed to assess the aspects of data sharing which researchers find to be most challenging (Stuart et al., 2017). In this survey, 63% of respondents reported that they shared data supporting their peer reviewed publications. However, researchers stated that their greatest barrier to data sharing is their ability to “organise data in a presentable and useful way”. A lack of time, concerns about copyright and licensing, and a lack of research funding for data sharing were also identified as practical challenges to increased data sharing in the survey.

Surveys from the publishers Wiley, Elsevier, and the publishing technology company Digital Science have found similar results regarding the proportion of researchers who share data (around two-thirds), and the ways by which researchers share data. The most common ways of sharing data that were reported tend to be suboptimal. All four surveys found a relatively low rate of repository usage by researchers, at 25% (Market Research, Wiley, 2017), (Berghmans et al, 2017), (Digital Science et al, 2017).

Although publisher policies and those of other stakeholders such as funders generally encourage data sharing using repositories, it is apparent that researchers do not feel adequately equipped to organise and describe their research data. One of the conclusions of the Springer Nature survey was that researchers should have faster and easier routes to data deposit, which do not require them to become experts in data curation. It also suggested that close collaboration between stakeholders, including researchers, research infrastructure providers, institutions and publishers will be necessary to affect change, and to develop solutions which simplify workflows for data deposit.

## Development of a third-party curation service

To help address the lack of time, skills or expertise needed to organise and share data reported by researchers in the survey, the Springer Nature data publishing team began the development of a curation service aimed to provide researchers with a means to deposit their data in a suitable repository, without requiring them to learn the skills necessary to create high quality metadata records. Such a service has a number of requirements, including a portal where data can be uploaded, and a platform (a repository) where the data can be assigned a DOI and published. The service also needed to include appropriate editorial checks to ensure data types that need to be deposited in discipline-specific repositories, such as DNA/RNA sequence data are directed to community specific repositories rather than a general repository, upon which the Research Data Support service is based. Additionally the service needed to ensure published data are based on scholarly research, and that sensitive data and data derived from human participants are suitably anonymised.

While the technical infrastructure and initial screening of submission was an important concern, a key focus of the service development was on drafting complete, accurate and appropriate metadata on behalf of the researcher. The intention was also to align the metadata being created with the FAIR Data Principles (Wilkinson et al, 2016), guaranteeing that any researcher who published data through the service could be assured that their data would be, at a minimum, Findable and Accessible. The metadata records created needed to be of high quality, and to add value to the data in a way that a researcher using the service could not have achieved themselves without specialist data curation skills.

Many repositories, and the documentation relating to standard metadata schema, provide guidance on the content and completeness that is expected when publishing data. The FAIR Data Principles also provide some high-level guidance on what can be expected of Findable and Accessible datasets, for example that their metadata includes a persistent identifier and a rich description. More recently the Go Fair website has expanded on what the term rich metadata implies, noting that it should be “generous and extensive, including descriptive information about the context, quality and condition, or characteristics of the data.”^2^ In spite of existing guidance on how high quality metadata should be created, there is a lack of documentation on how metadata quality can be assessed or benchmarked after the metadata have been created. Furthermore, the focus of the Springer Nature Research Data Support service is on curating datasets that support peer-reviewed publications, taking into account the needs of journal editors, peer reviewers, readers, as well as authors (data creators).

As we began to develop standard workflows and metadata descriptions for the new curation service, it became clear that an objective assessment would be necessary to allow the quality of the metadata created by the service to be quantified, and for the editorial standards that had been developed for the curation service to be further tested.

## Assessing metadata quality: methodology

Early in the development of the data curation service, a small group of stakeholders came together at an internal workshop to assess the ways that journal authors currently share data, and to try to objectively quantify the quality of associated metadata using a short survey form. This pilot workshop was held in 2016, and informed the more comprehensive metadata quality assessment which is described in detail here. In February 2017, the second workshop was held, with the aim of undertaking a comparative assessment between metadata already published in generalist repositories, and metadata which had been created according to the standards developed by the Research Data Editors, primarily Graham Smith.

In preparation for the workshop, 20 datasets that support peer-reviewed publications were identified by searching for repository names and Digital Object Identifier (DOI) prefixes in journals on the BMC, SpringerLink, Nature and EU PubMed Central websites. As the Springer Nature curation service workflow planned to deposit into the generalist repository figshare, the search was limited to datasets which were available in generalist repositories including figshare and Zenodo, and those which had been published using the Electronic Lab Notebook platform LabArchives, which also enables its users to publish datasets and assign them DOIs. Datasets published in the generalist repository Dryad were excluded from the search as these are subject to basic curation as standard by the repository’s curators. Data considered out of scope for the service include types that belong in structured, discipline-specific repositories, where a clear community mandate or expectation for deposition in these repositories exists. To curate data types the service would most likely receive, simple stratified sampling was performed to return:

- 50% tabular data (e.g. Excel, comma-separated text, SPSS or Stata files
- 25% code (e.g. R, PERL, Python files)
- 25% ‘other’ data encompassing media, bioinformatics and GIS format (e.g. waveform, video, image, Nexus and ArcGIS files)

A total of 50 datasets were returned as the initial sample. Four were subsequently removed from consideration as they also contained files that belonged in a specialist community repository. Twenty of the remaining 46 datasets were then randomly selected for curation. Once identified, the sample datasets and their metadata were downloaded by Graham Smith. In line with the original sampling, the datasets represented a range of disciplines, and the file formats included software code, spreadsheets, imaging and audio files.

The sample datasets were accompanied by varying levels of metadata, depending on the repository where they had been made available and the amount of information which the depositor chose to include. Most included the depositor’s name and a licence, accompanied by descriptive metadata which ranged from nothing at all, to one line, to a number of paragraphs. In some cases keywords had also been added to the metadata record.

These sample datasets were prepared to allow a participant-blinded comparison between metadata which had been prepared by researchers themselves, and metadata which had been curated by Springer Nature Research Data Editors. The downloaded datasets were curated by the Research Data Editor Graham Smith, using the standard process for metadata checks and enhancements which had been created during the development of the planned third-party curation service. This standard process creates metadata in a Dublin-core compliant format, and provides the Research Data Editor with guidance on filling out each metadata element. The information used to create the descriptive metadata is elicited from the dataset itself and from the manuscript or the published article associated with it. Curation in this context does not involve direct edits to the data files by the curators and is focused on the metadata only. The scope of the envisaged service did extend to providing advice to researchers on edits that could be made to data files, but this was not relevant to the assessment process outlined here. While the service is not intended to guarantee that data files are reusable, making data Findable and Accessible can lead to improvements in the reproducibility of the research. Spot checks are also carried out on the contents of files where these are accessible by the curator – for example to guard against the unintended publication of information about identifiable individuals.

Before beginning to add metadata to the data files as part of the standard curation process, they are split into appropriate collections and sub-collections if necessary. The descriptive metadata is intended to include contextual details relating to the research study which generated the data, and its methodology, as well as an overview of each file in the dataset. Additional information such as the software necessary to open particular file types may also be included if relevant. The metadata record is expected to act as a standalone description of the dataset, allowing the data to be interpreted without the need to read the associated research paper. In creating a description sufficient for this purpose but without replicating elements of the related manuscript, such as the methodology section, standardised levels of detail in the data description are required. This was a key challenge addressed by the assessment of the curated outputs of the service. Additionally, keywords and categories are included in the metadata to allow the dataset to be found more easily in the repository and via any data aggregators. The completed metadata record also includes a link to the DOI of the data’s associated peer-reviewed publication and clear citation information and a standard licence to support reuse of the data. While some additional functionality is provided by the figshare repository infrastructure used by the data publishing team (for example the ability to preview many data formats in the user’s browser, without the need to download them) the intention was to assess the quality and usefulness of the metadata provided, rather than the differences in repository features. An example of a published dataset which has been curated by the service (Rakotoniaina et al, 2017) is shown in Figure 1.

**Figure 1.**
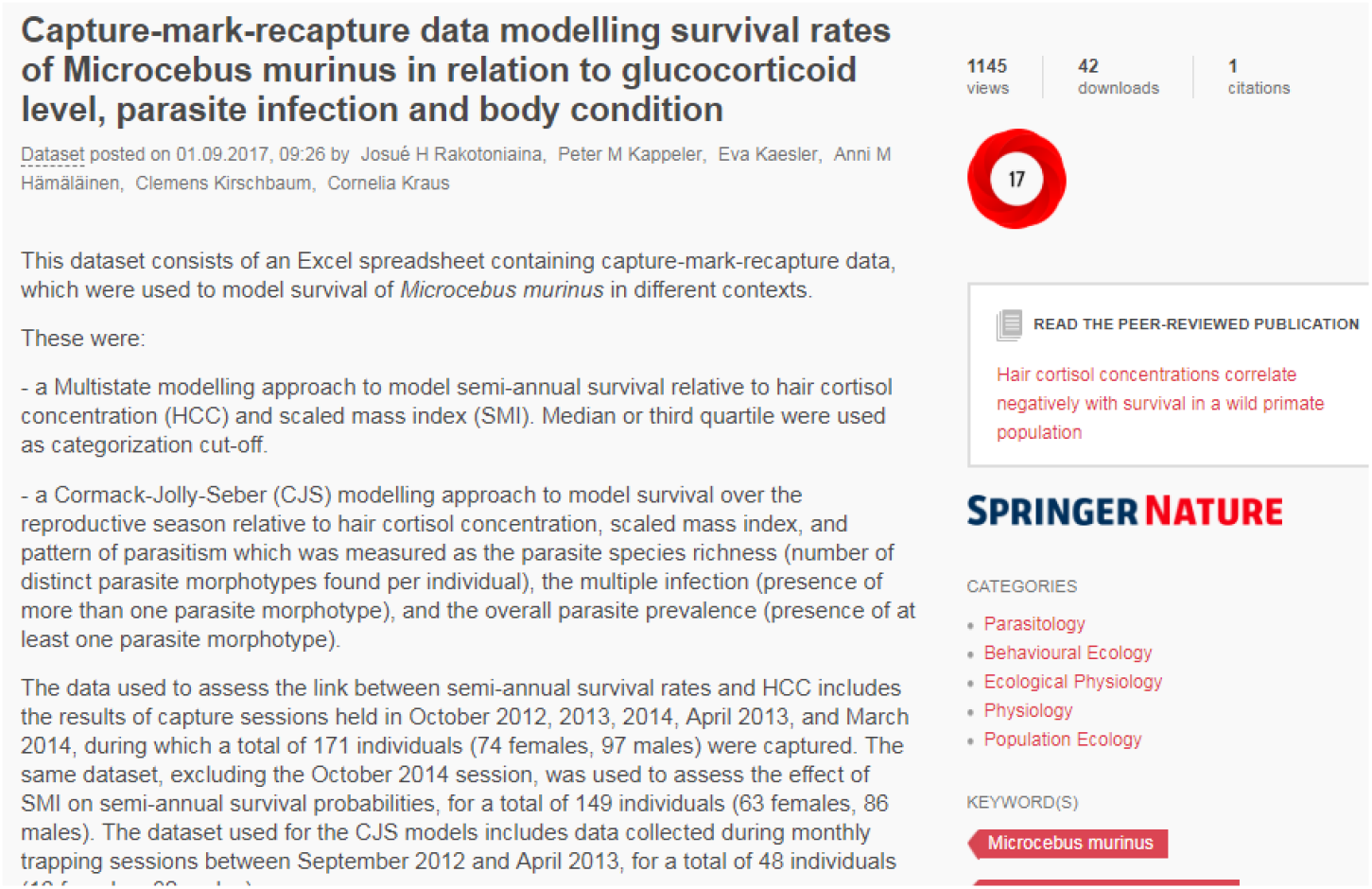
An example of a dataset which has been curated by the Springer Nature Research Data Editors.

Once the datasets had been curated using the standard process described above, they were made available in a private, unpublished (staging) area of the Springer Nature figshare repository. In addition to these 20 “edited” datasets, the original versions, not edited or enhanced by the Research Data Editors, remained in their original repositories where they were openly accessible. The sample data used during the workshop therefore consisted of 20 “edited” and 20 “unedited” datasets.

Ten professional Editors employed by Springer Nature (several of them former researchers) took part in the workshop, representing a range of disciplines (including genomics, neuroscience, physics and energy) and journals. During the workshop each Editor was randomly assigned two “edited” and two “unedited” datasets, and asked to review the quality of the metadata for each using a short survey form. The assignment process, while random, ensured that no editor assessed both the “unedited” and “edited” version of the same dataset. In an attempt to avoid potential bias, Editors were not informed that we would be comparing the quality of edited and unedited datasets, or that some of the datasets had been edited by the Research Data Editors.

The metadata quality survey (Smith, Hrynaszkiewicz & Grant, 2018) included questions regarding the clarity of the metadata, whether it was sufficient to allow a citation to be created, whether keywords had been included, if the file formats were clear and whether it included information that would be needed by another researcher in order to reuse the data. Likert scales were also used to allow the editors to quantify how complete the metadata record was, and its quality. Following the workshop the survey data were collated and a report was drafted by Graham Smith and lain Hrynaszkiewicz, and shared with the workshop participants and stakeholders involved in the development of the data curation service. An aggregated version of the survey responses was also published in the Springer Nature figshare repository.

## Data analysis

The data gathered during the workshop provided insight into the differences between the metadata records which had been published by researchers themselves, and those which had been edited using the metadata guidance developed by Springer Nature Research Data Editors. The survey responses allowed for precise analysis of how individual service features performed, as well as the value of the service. Overall, datasets which had been edited by the Research Data Editors scored more highly for quality and completeness, rating 4.1 out of 5.0 for overall quality, compared to 2.8 for unedited (Figure 2). Datasets which had not been edited were described, in comparison, as difficult to find or access, with metadata which was often missing, or of low quality.

**Figure 2.**
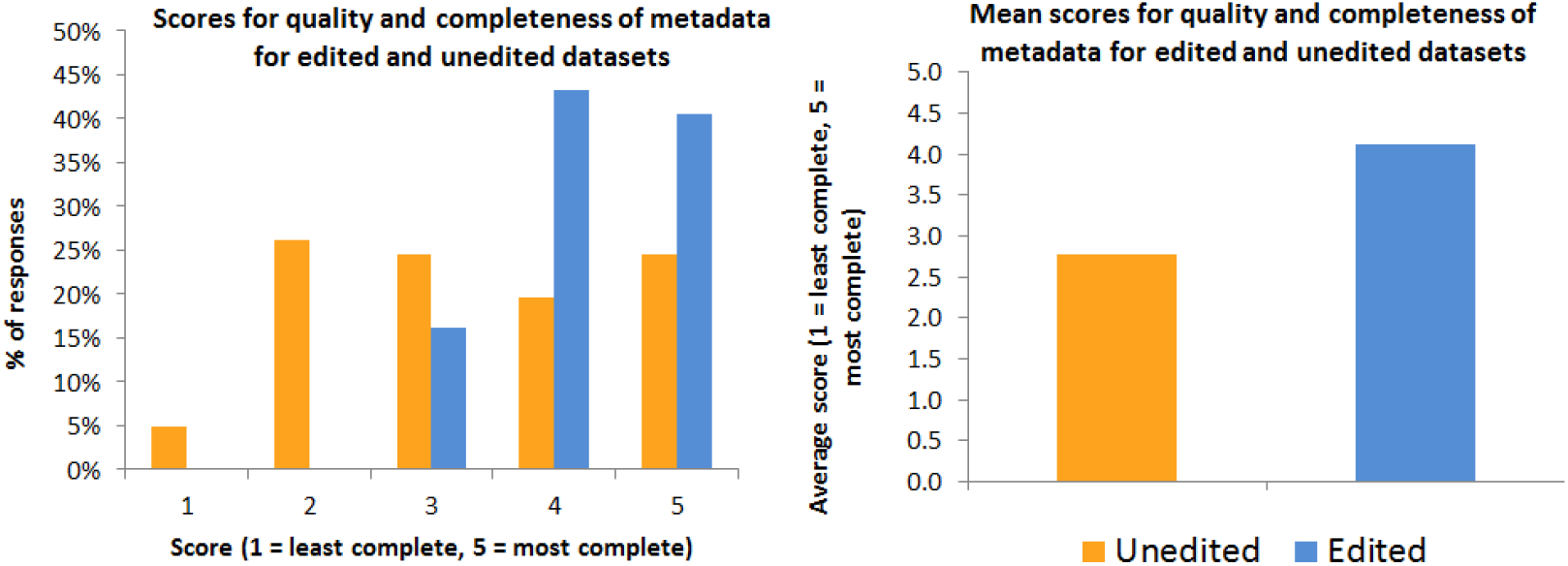
‘Quality and completeness’ scores for edited and unedited datasets.

Additionally, when asked whether it was clear what the data were and how they had been generated, 100% of the edited datasets were found to be clear while only 41% of the unedited datasets were found to provide the information clearly (Figure 3). The lack of clarity for the unedited datasets represents a barrier to reuse, as those accessing the datasets are unlikely to be able to assess whether the data are relevant or useful to them.

**Figure 3.**
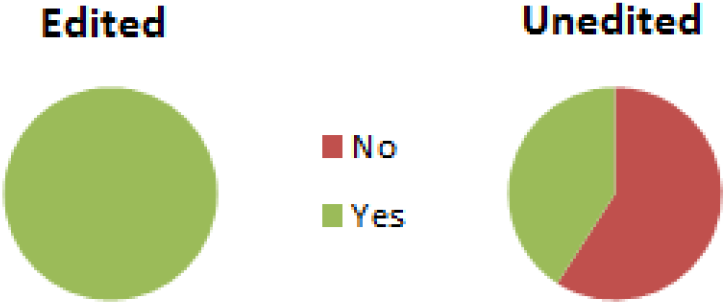
Responses to the question ‘Is it clear what kind of data they are, and how they were generated?’

Data were also gathered in relation to metadata completeness, as editors were asked whether the datasets they assessed had keywords included, and whether these were found to be useful. 100% of the edited datasets were found to have keywords, all of which were considered to be useful; in comparison, only 36% of the unedited datasets included keywords and of these, less than half (44%) were considered to be useful additions.

Editors were also asked to assess whether the datasets had clear terms of use included in their metadata, for example through the availability of a Creative Commons licence. 100% of the edited datasets had terms of use included, while 73% of the unedited datasets did. It is not evident whether the availability of terms of use for the unedited datasets was improved by mandatory requirements of the repositories where they were deposited, or whether researchers are more likely to understand the implications of adding a licence compared to the value of adding keywords.

Data relating to citations and links between datasets and publications were also gathered, as editors were asked whether they believed that a data citation (a formal bibliographic reference to the dataset) could be drafted based on the metadata which accompanied each dataset, and whether the paper which corresponded to the dataset was linked to it or could otherwise be easily found. The editors believed that 83% of the edited datasets could be used to create a data citation, while 73% of the unedited datasets had sufficient information included. The quality of links from the data to the associated papers also contrasted between the edited and unedited datasets, with 89% of the edited datasets having a paper which was easy to find using the dataset’s metadata, in comparison to only 36% of the unedited datasets (Figure 4).

**Figure 4.**
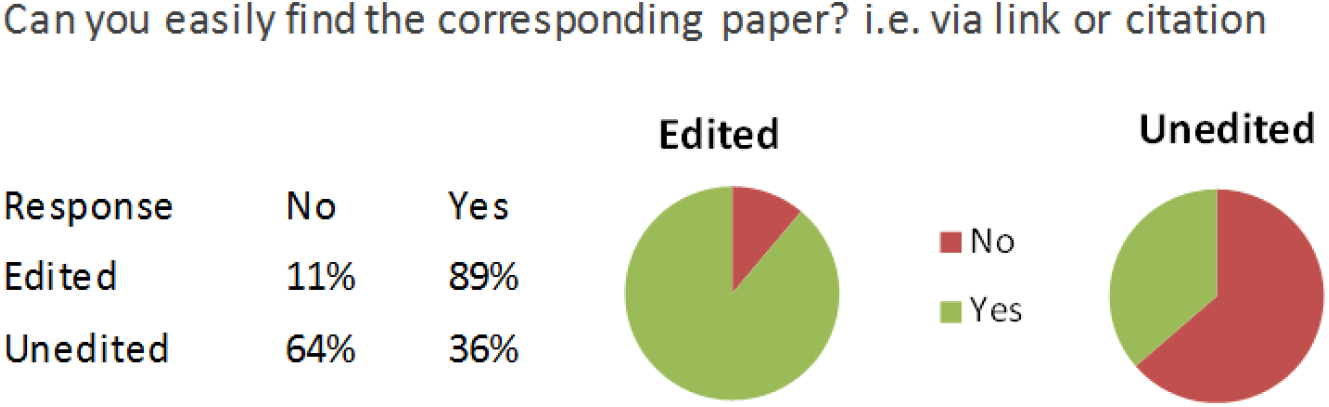
Responses regarding the availability of links between published datasets and their associated papers.

Note: a technical issue with data access led to 18 edited and 22 unedited datasets being assessed. Results are therefore provided in percentages to allow for direct comparison of the two groups.

Through the analysis of the data gathered during the workshop, it was evident that there was a discernible difference between the metadata provided for the edited and unedited datasets. The unedited datasets were associated with poorer quality metadata which was less complete, and therefore provided less information to those wishing to understand or reuse the data. The unedited datasets were also less likely to include useful keywords, which indicates that it would be more challenging for users to find and access these datasets in the first place. Additionally, it was more difficult for users to find the publication associated with unedited datasets, compounding the issue of poor descriptive metadata which could have been mitigated by accessing the publication and reading more about the methodology used for data generation.

As well as scoring highly in comparison to the unedited datasets, the edited datasets also scored objectively highly across all of the assessment questions. The consistent approach to metadata creation ensured that all of the edited datasets had a clear licence in place which could be easily found, and they all had useful keywords included. All of the edited datasets were also reported as having a clear description of what the data consisted of and how it had been generated. The analysis of the assessment of the edited datasets indicated that the metadata guidelines used to describe the data were aligned with the intention to ensure that the datasets would be both findable and accessible. Editors also provided qualitative feedback in the form of additional comments relating to each dataset. Specific comments were used to refine the curation standards, for example the structure of the data description, the value of including elements from the related publication, explanations of discipline-specific abbreviations and what files would be expected in a data record versus the publication’s supplementary information.

## Limitations

Although the intention of the metadata assessment was to allow an analysis of the quality of a set of metadata records, the subjective viewpoints of the editors mean that the assessment cannot be considered truly objective. Additionally, although the purpose was not to assess the functionality or user-interfaces of the repository infrastructures examined, the features or limitations of each repository may have impacted positively or negatively on the scores for each dataset depending on where they had been published. Given the broad range of data types and disciplines that are in scope for Research Data Support, a wider range of data records and Editors would be required to make a similar assessment process truly comprehensive. Furthermore, as a single-blind study in which the experimenters were not blinded to which datasets had been edited and which had not been edited, the observer-expectancy effect should also be considered.

## Conclusion

Following the metadata quality assessment workshop, the development of the third-party curation services continued, and launched under the name Springer Nature Research Data Support in March 2018 following a small scale pilot in 2017. The analysis of the data gathered during the metadata quality assessment contributed to shaping the standards and workflows which are now in place as part of the service. The metadata creation process for the service includes the drafting of a comprehensive description to accompany each dataset; the addition of keywords; inclusion of creators and funders associated with the dataset; checks and advice on the licences which have been applied; and the creation of a data citation and data availability statement to be added to any manuscript which relates to the dataset.

The service has been made available to authors through the manuscript submission system at certain Springer Nature journals, as well as being provided to any researcher who has already published an article with Springer Nature or any other trusted publisher who has data which they would like to share. The services have also been provided to conference attendees to support the publication of their data alongside conference proceedings. Over one hundred datasets have now been published by the services, with datasets accompanying articles in Springer Nature journals including *Nature* (Giles, Xu, Near & Friedman, 2017) and *BMC Ecology* (Rakotoniaina et al, 2017), and to accompany articles published by other publishers such as the Journal of Experimental Biology (Jacobs & Holzman, 2018).

The workshop also provided the authors with quantitative data regarding the improvements that the service made to the datasets which were analysed. Specific feedback from editors provided a basis from which to refine our curation standards, for example on the level of description required around methodology or code operation for software datasets, and metadata that was more relevant to the related publication than the dataset itself.

Since the launch of the curation service, the Research Data Editors who form the curation team have continued to develop the guidelines which inform how metadata should be created. As new data types are submitted for curation, additional use-cases have been addressed, for example approaches to describing complex datasets created by computer science researchers which consist of hundreds of individual files. More consistent approaches to language and terminology are also being formalised to ensure that aspects of the description such as sample size are described in a consistent way. The feedback gained from this assessment by professional Editors helped to shape the standardisation of key elements of a detailed data description. Although the metadata assessment exercise has not been repeated, researcher surveys and interviews have been undertaken to gather feedback from those using the curation service. Additional work is also underway to expand the scope of the metadata curation to include data which are too sensitive to be shared openly, and data which are not yet associated with a manuscript or a published research article.

While assessment of metadata quality is just one measure of the impact of a third-party data curation service, quantifying the value added by the service is important in considering its sustainability and future business models. Future work to assess the value added by curation could consider the usage of curated datasets compared to non-curated datasets, such as through analysis of downloads, views and citations. Anecdotally, a published peer reviewer report about an article that included data curated using the Research Data Support service noted that “supporting data are very complete, including experimental procedure” (Liang, 2018). Assessing the benefits of curation for supporting the peer review and editorial process of journals is another potential avenue of further research. Markowetz has claimed that making data available clearly and transparently enables papers to be published more quickly, as editors and reviewers can more efficiently assess the authors’ methodology (Markowetz, 2015). Other groups are also developing tools, for example, to assess “fitness” of data for reuse.^3^

Before the metadata assessment was undertaken, there was no standard methodology available to assess or benchmark research data curation quality. The survey questions which were used during the workshop are therefore available to other metadata creators who wish to undertake a similar assessment of their metadata quality.

## Data availability

An aggregated version of the survey responses, including the survey questions used, is available in the Springer Nature figshare repository, accessible at https://doi.org/10.6084/m9.figshare.6200357.v1 (Smith, Hrynaszkiewicz & Grant, 2018).

## Acknowledgements

The authors would like to thank the editors who participated in the workshop for their contribution to this research, and Andrew Hufton for his assistance in developing the curation standards described.

## Footnotes

1 Data policy standardisation and implementation Interest Group: https://www.rd-alliance.org/groups/data-policy-standardisation-and-implementation

2 Go Fair: https://www.go-fair.org/fair-principles/f2-data-described-rich-metadata/

3 WDS/RDA Assessment of Data Fitness for Use Working Group: https://www.rd-alliance.org/groups/assessment-data-fitness-use

